# Ebselen Reacts with SARS Coronavirus-2 Main Protease Crystals

**DOI:** 10.1101/2020.08.10.244525

**Authors:** Tek Narsingh Malla, Suraj Pandey, Ishwor Poudyal, Denisse Feliz, Moraima Noda, George Phillips, Emina Stojkovic, Marius Schmidt

## Abstract

The SARS coronavirus 2 main protease 3CLpro tailor cuts various essential virus proteins out of long poly-protein translated from the virus RNA. If the 3CLpro is inhibited, the functional virus proteins cannot form and the virus cannot replicate and assemble. Any compound that inhibits the 3CLpro is therefore a potential drug to end the pandemic. Here we show that the diffraction power of 3CLpro crystals is effectively destroyed by Ebselen. It appears that Ebselen may be a widely available, relatively cost effective way to eliminate the SARS coronavirus 2.

## Introduction

During the COVID-19 pandemic in the last half year, a large body of literature became available to understand and combat the SARS coronavirus-2 (CoV-2). Due to its importance for virus protein maturation the SARS CoV-2 main protease (Mpro), also called 3CLpro, became a major drug target. In its functional form the 3CLpro is a homo dimer with His/Cys dyads (His-41 and Cys-145) in the active centers^1^. The structure of the CoV-2 3CLpro has been solved recently (Fig. 1) guided by high similarity to other coronavirus 3CLpros^1^. A large number (>500) of SARS CoV-2 3CLpro structures were recently deposited in the protein data bank mainly following a fragment screening study at the synchrotron Diamond near Oxford, England. Binding of fragments not only to the active center was observed. They all constitute a database of potential drugs to target the 3CLpro. Apart from the fragments, the most promising compounds are the α-ketoamides (Fig. 1) which bind tightly to the 3CLpro^1-3^. As they are complicated to synthesize they carry a hefty price tag (companies ask for 14,000 dollars/g as recently inquired by one of the authors). A less expensive, but also less known compound that binds to the 3CLpro is Ebselen^4^. Ebselen is a selenium compound (Scheme 1) currently tested for a number of diseases such as bipolar disorder and hearing loss^4^. Selenium is an essential metal, but toxic in higher doses. Ebselen has been shown to bind strongly to the CoV-2 3CLpro^4^, but the structure of the complex is unknown. Here we show what happens when Ebselen is added to 3CLpro crystals.

**Figure 1.**
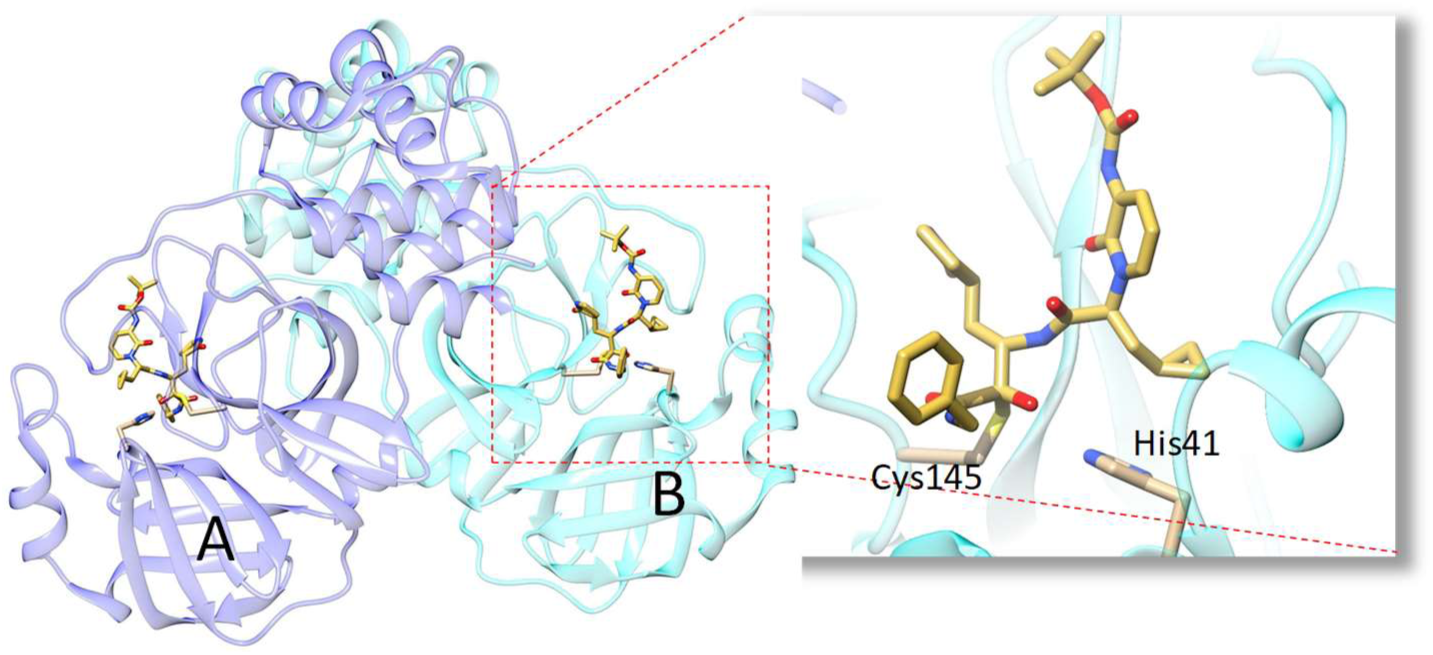
SARS CoV-2 3CLpro in its functional, dimeric form^1^. The two subunits A and B are shown in dark and light blue. An α-ketoamide inhibitor is bound to the active site (red box, enlarged). The His-41/Cys-145 catalytic dyad is marked.

## Methods

### Expression

**Scheme 1.**
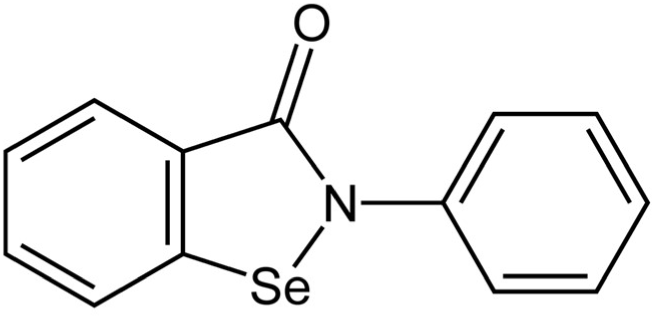
Chemical structure of Ebselen, an organo-metallic compound.

The CoV-2 3CLpro sequence was synthetized (GenScript) for optimized expression in E. coli according to sequence information published previously^1^. In short, the N-terminus of 3CLpro is fused to glutathione-S-transferase (GST). It further has a 6-His tag at the c-terminus. The N-terminal GST will be autocatalytically cleaved off after expression due to an engineered 3CLpro cleavage sequence. The His tag can cleaved off by a PreCission Protease. Overexpression and protein purification protocols were modified from previous reports. E. coli were grown to 0.8 OD600 at 37oin terrific broth. Expression was induced by 1 mmol/L IPTG at 25° C. After 3 h of expression, the culture was induced a second time (1 mmol/L IPTG), and shaken overnight at 20° C. The yield is about 80 mg for a 6 L culture. Cells were resuspended in lysis buffer (20mM Tris Base, 150 mmol/L NaCl, pH 7.8.). After lysis of the bacterial cells, debris was centrifuged at 50,000 g for 1 hour. The lysate was let stand at room temperature for at least 3 h (overnight is also possible). After this, the lysate was pumped through a column containing 15 mL of Talon Cobalt resin (TAKARA). The resin was washed without using imidazole using a wash cycle consisting of low salt (20 mmol/L Tris Base, 50 mmol/L NaCl, pH 7.8), high salt (20 mmol/L Tris Base, 1 mol/L NaCl, pH 7.8) and low salt (as above) solutions (about 20 column volumes each). After the wash cycle was completed, the column was let stand for an additional 2 h at room temperature followed by another wash cycle. The final product was eluted by 300 mmol/L imidazole, dialyzed immediately in 20 mmol/L Tris base, 150 mmol/L NaCl, 0.1 mmol/L dithiotreitol (DTT), pH 7.8, and concentrated to 20 mg/mL. Note, the c-terminal 6-His tag was not cleaved off. Due to this one step purification protocol, only 24 h are required from cell lysis to the pure 3CLpro product. The product is within 1.7 Da of the theoretical molecular weight as determined by mass spectroscopy.

**Figure 2.**
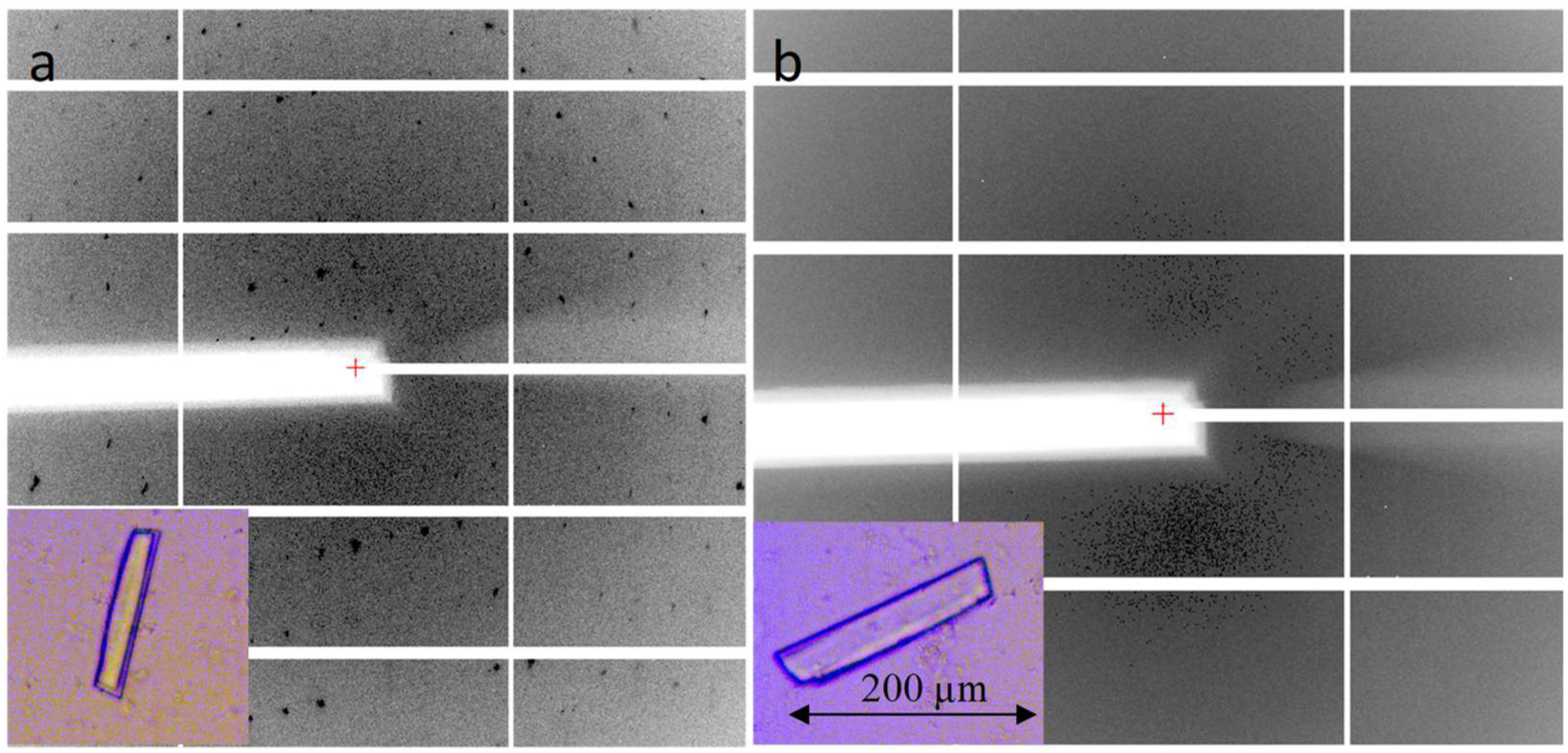
Low resolution (∼3.5 Å in the corner) diffraction patterns of SARS CoV-2 3CLpro. (a) no ligand, (b) with Ebselen. Inserts: crystal images w/o and w Ebselen, respectively

### Crystallization

The concentrated 3CLpro (with the His-tag left on) was diluted to 4 mg/mL. 100 μL of the diluted 3CLpro was mixed (1:1) in batch mode with the same amount of 25 % PEG 3350, Bis-Tris 100 mmol/L, pH 6.5. A few days later, nicely shaped crystals with dimensions of about 200 × 30 x 30 μm^3^ were appearing. Crystals were soaked in mother liquor by adding Ebselen in powder form as Ebselen is not very soluble in water. Enough Ebselen that would otherwise produce a 50 mmol/L solution was added. After 2 days of soaking, the crystals completely maintained their morphology (Fig. 2b).

### Data Collection and Structure determination

The crystals were mounted in Mitegen microloops (30 - 50 μm) and directly frozen in pucks suspended in liquid nitrogen for automated (robotic) data collection. The dewar with the pucks were driven to the Advanced Photon Source for robotic data collection at Sector-19 (Structural Biology Center, SBC, beamline 19-ID-D). Data collection was fully remote due to restriction of the COVID-19 pandemic. Fig. 2 shows a comparison of diffraction patterns collected from untreated crystals (Fig. 2a), and crystals treated with the Ebselen (Fig. 2b). As the untreated crystals diffracted beyond 2 Å, the treated crystals did not show any Bragg reflections whatsoever, even at lowest resolution. They are completely amorphous, despite the nice crystal-like shape. Accordingly, the structure of only the untreated 3CLpro can be solved. A dataset to 2.2 Å was collected (0.5° rotation and 0.3 s exposure per detector readout for a total of 180°). Data was processed with HKL3000^5^. Data statistics in shown in Tab. 1. Initially the spacegroup was found to be C2 (monoclinic centered). It was further determined by the CCP4 program^6^ *pointless* that spacegroup I2 (monoclinic body centered) is more suitable, in accordance with published results^7^. The 3CLpro structure with pdb access code 6WQF^7^ was used as initial model. Molecular replacement was not necessary. The model fits immediately and can be used for refinement. Refinement was done using refmac^8^ (version 5.8.0238). The structure is shown in Fig.3. Its mean square deviation from model 6WQF is about 0.4 Å with 0.5 Å standard deviation. This means that the 3CLpro cryo structure with the His-tag and the room temperature structure are identical within the resolution limit. The structure of the His-tag could not be determined. There is, however, a large crystal cavity near the C-terminus that might accommodate (a very disordered) 6 histidine structure.

**Table 1.** Data collection and refinement statistics*

|  | SARS CoV-2 3CLpro |
| --- | --- |
| Beamline | 19-ID-D, APS |
| Resolution | 2.2 Å |
| Temperature | 110 K |
| Space group | I2 |
| Unit-cell parameters (°,) | a = 44.7 Å b = 53.4 Å c= 112.3 Å<br>$\alpha=90^\circ$ $\beta=100.0^\circ$ $\gamma=90^\circ$ |
| No of unique reflections | 9426 |
| Redundancy | 3.1 (1.9) |
| Completeness (%) | 70.0 |
| CC1/2 (%) | 99.8 (47.6) |
| R<sub>merge</sub> (%) | 7.3 (76.5) |
| Refinement |  |
| R<sub>cryst</sub>/R<sub>free</sub> (%) | 19.1/26.3 |
| RMSD to 6WQF<sup>7</sup> | 0.39 +/- 0.53 Å |

\*last resolution shell in brackets

**Figure 3.**
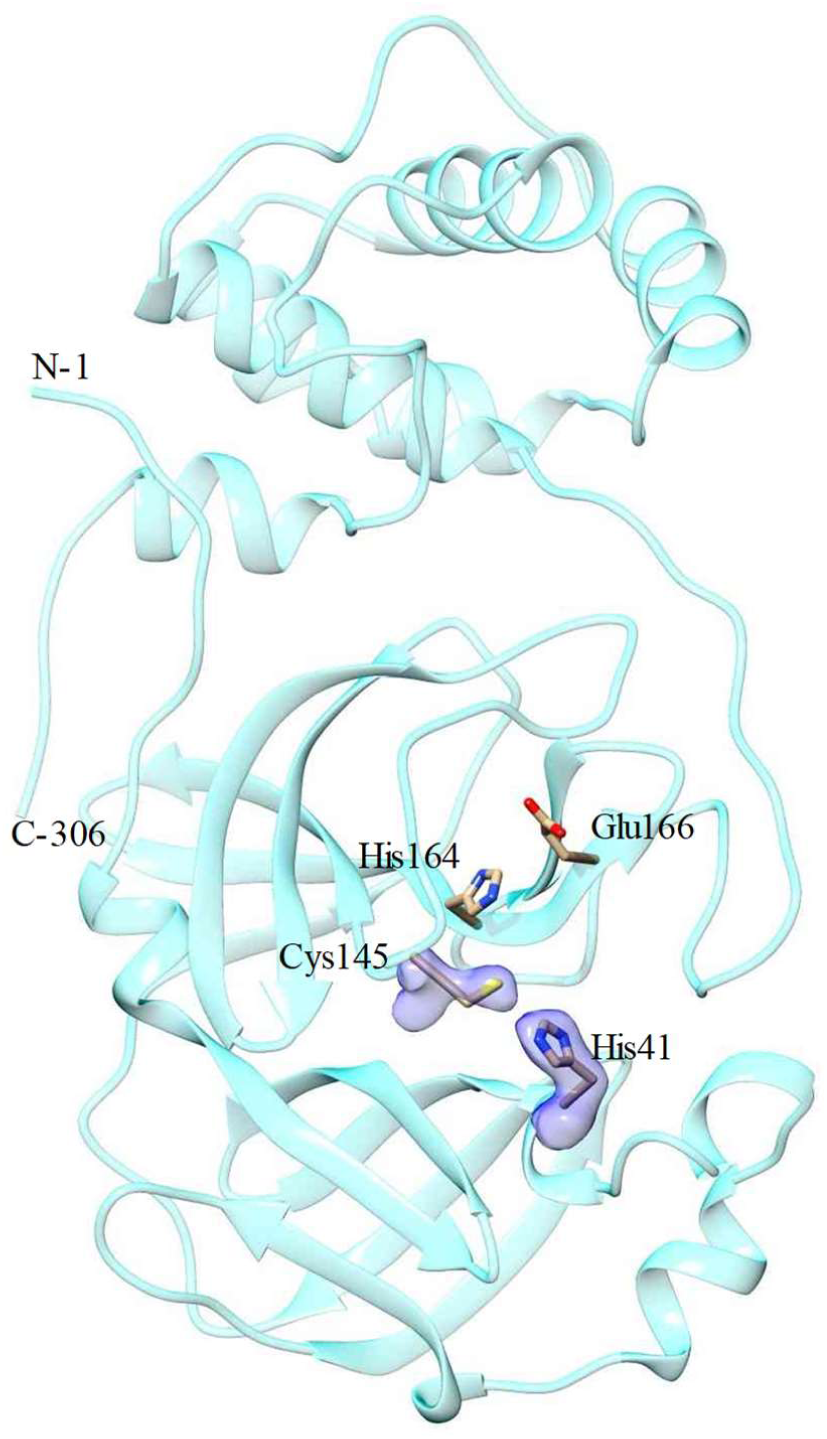
Structure of the SARS CoV-2 3CLpro without Ebselen as determined here. Electron density (1.2 sigma) is shown for the catalytic dyad. Other active site residues are also marked.

## Results and Discussion

A very important result is that c-terminal PreCission His-tag cleavage is not required to obtain 3CLpro crystals. This speeds up purification to the extent that now gram sized 3CLpro preparations can be obtained in a relatively short period of time. This is a prerequisite for mix-and-inject approaches at free electron lasers with gas-dynamic-virtual-nozzle type mixing injectors^9,10^. At XFELs low concentrations of Ebselen can be mixed with the crystalline slurry which is quickly injected into the X-ray beam. There is hope that crystal decomposition is slower than the delay between mixing and injection. Then, the structure of the Ebselen-3CLpro complex may be determined. In addition, microcrystals also tend to be unusually flexible and may survive large unit cell changes^11^, which might make these experiments even more conceivable.

Since Ebselen has this strong effect on 3CLpro crystals, it a clear indication for the tight affinity of this compound to the 3CLpro. Binding of Ebselen to the catalytic dyad has been already identified by mass-spectroscopy and published earlier^4^. However, also after the experiments reported here, the 3CLpro-Ebselen structure remains elusive. As Ebselen is a potent drug (and it is not too toxic even in higher concentrations^4^), it has to be seen whether it can be used as an effective weapon against the SARS CoV-2, and maybe all other pathogenic coronaviruses. Whether it is suitable for a drug must be determined by physicians in a clinical setting perhaps in combination with a SARS CoV-2 RNA dependent RNA polymerase inhibitor such as remdesivir^12^. As crystallographers and structural biologists, though, we can deliver evidence for a potential mechanism as shown recently by others for the α-ketoamides^1,2^ and also remdesivir^12^.

This manuscript is published without delay on BioRxiv to disseminate rapid methods for preparation, purification and structure determination of 3CLpro to a wide community. In addition, this manuscript intends to alert and increase awareness of the potential of Ebselen as a 3CLpro inhibitor and its value perhaps in combination, as the authors are not aware that anything is reported about it in the public press. The coordinates of the 100 K 3CLpro structure as determined here are available on request.

## Acknowledgement

This work is supported by NSF grant STC 1727290 (BioXFEL) and NSF grant RAPID 2030466. Results shown in this report are derived from work performed at Argnonne National Laboratory (ANL), Structural Biology Center (SBC) at the Advanced Photon Source (APS), under U.S. Department of Energy, Office of Biological and Environmental Research contract DE-AC02-06CH11357. The authors thank Darren A. Sherrell for setting up the experiment at the beamline.

